# Detection and quantification of single mRNA dynamics with the Riboglow fluorescent RNA tag

**DOI:** 10.1101/701649

**Authors:** Esther Braselmann, Timothy J. Stasevich, Kenneth Lyon, Robert T. Batey, Amy E. Palmer

**Affiliations:** Department of Biochemistry, University of Colorado, Boulder, CO, USA, 80303; BioFrontiers Institute, University of Colorado, Boulder, CO, 80303; Department of Biochemistry and Molecular Biology, Colorado State University, Fort Collins, CO, 80523; World Research Hub Initiative, Institute of Innovative Research, Tokyo Institute of Technology, Yokohama, 226-8503, Japan

## Abstract

Labeling and tracking biomolecules with fluorescent probes on the single molecule level enables quantitative insights into their dynamics in living cells. We previously developed Riboglow, a platform to label RNAs in live mammalian cells, consisting of a short RNA tag and a small organic probe that increases fluorescence upon binding RNA. Here, we demonstrate that Riboglow is capable of detecting and tracking single RNA molecules. We benchmark RNA tracking by comparing results with the established MS2 RNA tagging system. To demonstrate versatility of Riboglow, we assay translation on the single molecule level, where the translated mRNA is tagged with Riboglow and the nascent polypeptide is labeled with a fluorescent antibody. The growing effort to investigate RNA biology on the single molecule level requires sophisticated and diverse fluorescent probes for multiplexed, multi-color labeling of biomolecules of interest, and we present Riboglow as a new member in this toolbox.

## Introduction

Approaches to label and track cellular biomolecules using fluorescent tags have revolutionized our ability to investigate how their spatiotemporal dynamics relate to function in live cells^1^. While tagging proteins with fluorescent markers offers a versatile repertoire of choices in terms of color, labeling modality and detection mode^2^, labeling RNAs with genetically encoded tags has lagged behind due to the absence of naturally occurring fluorescent RNAs. Seminal work by the Singer lab^3,4^ established the MS2 system as a powerful tool to interrogate dynamics of mRNAs in live cells. The MS2 system typically consists of 24 stem-loop (SL) repeats genetically fused to a gene of interest in the 3’ untranslated region (UTR)^3,5,6^. Each SL binds a dimer of the bacteriophage-derived MS2 coat protein, which is genetically fused to GFP^3^. By tagging each RNA with 24 copies of the SL, up to 48 copies of MS2-GFP accumulate on each RNA molecule. The MS2 system has been instrumental for elucidating mRNA dynamics, and is the only RNA-tagging platform shown to enable single molecule RNA imaging in live mammalian cells. Here, we explore Riboglow as an alternative technique to label and track single RNAs in live cells in order to complement and broaden the scope of MS2-based single RNA tracking.

Visualizing mRNA with the MS2 system has enabled tracking of single mRNAs, characterization of their movement, and has revealed fundamental features of mRNA biology that cannot be studied without single molecule resolution. Tracking of single mRNAs revealed different types of movement (diffusive, corralled and directed)^3^ (see also Supplementary Table 1, Supplementary Fig. 1 for a summary), and tracking β-actin mRNA (encoded by ACTB) emerged as a model mRNA for single particle studies^7–13^. mRNAs have “zipcode” sequences, often in their 3’ UTR^14^. Interactions of these zipcode sequences with RNA binding proteins enables directed movement along the cytoskeleton^6^. Transport along the cytoskeleton, along with other mechanisms to distribute mRNA throughout the cell, results in distinct mRNA localization patterns in cells to ensure that the message is positioned close to where the translated protein is needed^15^. Single mRNAs have been tracked in whole organisms, including in yeast^4^, *Escherichia coli*^16^, and *Drosophila melanogaster*^17^ and the creation of a transgenic mouse^11,18^ allowed mRNA tracking in the whole mouse and in diverse primary cell types, including neurons^19^. Interestingly, different cell types such as neurons with protrusions exceeding hundreds of µm, and compact cell types such as fibroblasts, utilize different mechanisms to distribute β-actin mRNA^20^. The diversity of mRNA dynamics highlights the need to investigate mRNA movement in different cell types and interrogate the complex network of mRNA binding proteins and their role for distributing mRNAs within cells^21^.

Recently, the MS2 system was combined with antibody-based probes to gain insights in the mRNA life cycle. Translation of single mRNAs can be monitored by following the nascent peptide chain via a loaded affinity probe^22^ or the SunTag^23–25^; these studies from different labs yielded comparable protein translation kinetics (see ref^26^ for a comparison). Other aspects of the mRNA life cycle, such as interactions with stress granules and P bodies^27^, association with the endoplasmic reticulum^28^, mRNA frame shifting^29^ and nonsense-mediated mRNA decay^30^, were also characterized using a combination of MS2-based mRNA labeling and complementary fluorescent probes on the single molecule level. Flexibility of mRNA tagging with different fluorophores and labeling systems will be critical for combining different fluorescent probes and labeling systems in the future.

While the MS2 system has been critical for single mRNA tracking, its large size, the need for genetic incorporation of both the tag and the MS2-fluorescent fusion protein, and background signal from the unbound fluorescent protein represent limitations that have sparked efforts to develop additional RNA tracking platforms. Advances of the MS2 platform itself include development of the orthogonal PP7 system^31^, a split fluorescent protein system^32^, use of HaloTag^23^ to reduce background fluorescence, and engineering efforts to avoid MS2-tag induced changes in mRNA degradation^33^. More recently developed techniques to label RNA use smaller aptamer tags that bind small molecules, including the Spinach/Broccoli^34,35^ and Mango tags^36,37^, but single molecule detection in living cells has not yet been achieved with any small molecule-based approach. Entirely orthogonal tools have been developed in which fluorescent probes are delivered into cells to bind untagged RNA without the need to genetically add tags^38–40^, enabling in principle visualization of any endogenous RNA of any size, such as miRNA^41,42^. Finally, additional approaches for tracking RNAs include incorporation of fluorescent nucleotide analogs to study mRNAs encoding mitochondrial proteins translated on endosome-containing ‘hotspots’ in axons^43^, molecular beacons for studying co-transcriptional vs. post-transcriptional alternative splicing^44^, and micro-injected fluorescent RNA probes to examine the miRNA life cycle^41,42^. While delivery of probes into cells may be technically challenging, advances to improve delivery routes of various fluorescent molecules in live cells have been explored^26,45,46^. Despite these advances, the growing field of RNA biology is in need of diverse and robust fluorescence tools to visualize RNAs with single molecule detection in live cells to gain insights in the complexity and regulation of RNA dynamics.

We recently developed Riboglow, a platform to tag RNAs with a short aptamer tag that binds a small molecule and induces fluorescence increase^47^. Riboglow relies on the bacterial cobalamin riboswitch, which binds cobalamin with a nM K_d_, genetically fused to an RNA of interest. Cobalamin has fluorescence quenching properties, and we synthesized and validated a suite of cobalamin-fluorophore probes where fluorescence is quenched relative to the free fluorophore. The RNA tag binds cobalamin, inducing a conformational change and hence an increase in fluorescence. This architecture allows us to easily pick and choose the organic probe moiety and thereby color. We demonstrated recruitment of mRNAs to stress granules and tagging of the short non-coding U1 snRNA using this platform^47^. Here, we extend this work and demonstrate tracking of single mRNAs with Riboglow. The Riboglow platform promises to be a powerful addition to the growing toolbox for live cell RNA imaging, especially when single molecule sensitivity is possible.

## Results

To assess if single mRNAs can be detected and tracked with our Riboglow platform^47^ in live cells, we tagged ACTB mRNA with 12 copies of the Riboglow RNA tag (Supplementary Figure 2). We chose to visualize ACTB mRNA because single ACTB mRNAs have been detected and tracked with the MS2 system in a variety of cell types and experimental conditions^3,7–12,48^. This robust body of existing studies enables us to benchmark detection sensitivity and results from Riboglow tagging (see Supplementary Table 1 and Supplementary Figure 1 for an overview of literature data and experimental conditions). We chose Cbl-4xGly-ATTO 590 as the fluorescent probe for Riboglow, as this probe exhibits ∼5-fold fluorescence turn-on upon binding to the RNA tag and minimal photo-bleaching^47^. Cells were imaged with highly inclined and laminated optical sheet (HILO) microscopy^49,50^, an image acquisition modality commonly used for single particle tracking in live cells that allows single particle detection throughout the cell volume while minimizing photo-bleaching^22,42,49,51^. We readily detected distinct puncta in the ATTO 590 channel (Fig. 1A) that moved rapidly throughout the cell when a plasmid to produce the Riboglow-tagged ACTB was transfected (Supplementary Movie 1). Fluorescent puncta were detected over background using the FIJI plugin Trackmate^52^ (Fig. 1B), enabling us to develop a detection and tracking pipeline for these particles (see Methods for parameters used for single particle detection).

**Figure 1:**
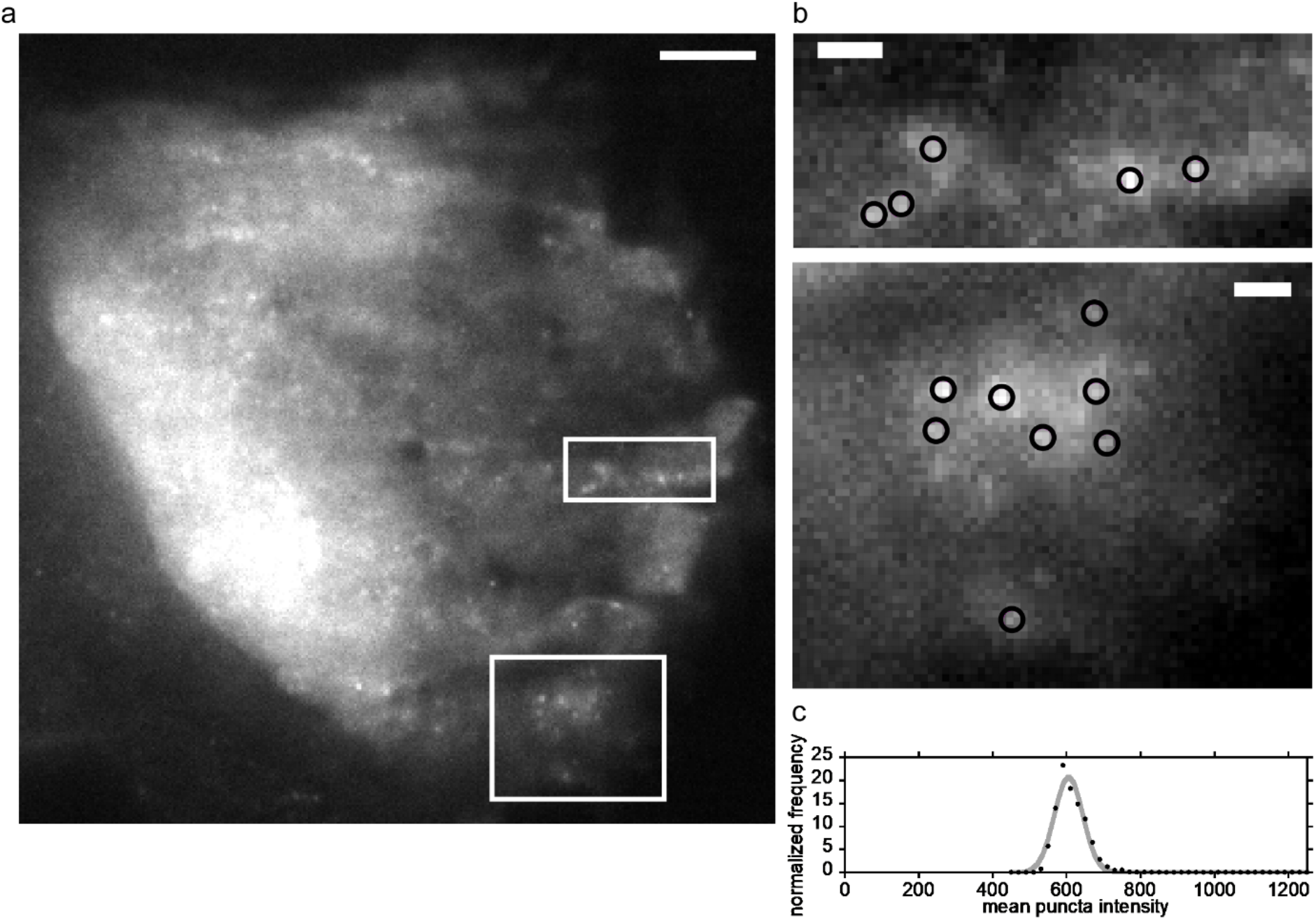
Detection of single mRNAs using the Riboglow platform. (A) U2-OS cells were transfected with a plasmid encoding ACTB mRNA, tagged with 12 copies of Riboglow. Twenty-four hours after transfection, Cbl-4xGly-ATTO 590 was bead loaded and cells were imaged live. Fluorescent puncta were readily detected in the ATTO 590 fluorescence channel. Scale bar = 5 µm. (B) Zoomed in view of areas indicated by white boxes in (A). Scale bar = 1 µm. Black circles indicate particles detected with the FIJI tracking plugin TrackMate^52^. (C) Intensity distribution of the mean puncta intensity for particles detected for the entire image acquisition for select ROIs. Data shown here were fit using a single Gaussian distribution. n = 2764 puncta, R = 0.98.

We determined that the observed puncta represent single mRNAs, based on the following observations. (i) When we compared cells with and without transfection of the Riboglow-tagged ACTB, no fluorescent puncta were seen in the untransfected control (Supplementary Movie 2 vs. 3). (ii) The distribution of intensities of individual puncta in each region of interest (ROI) analyzed by Trackmate follows a Gaussian distribution (Fig. 1C), as others observed for tracking of single MS2-tagged ACTB mRNA^7^. We reasoned that a Gaussian distribution is indicative of a single species of molecules, as one would expect for single molecules. In some ROIs, we detected two populations across the entire movie (Supplementary Fig. 3). To exclude potential artifacts, only ROIs with a single population were used for further analysis. (iii) We observed similar behavior for MS2-tagged ACTB mRNA in our hands, as follows. We transfected a construct producing ACTB tagged with 24 copies of the MS2-SL (Supplementary Fig. 2) into U2-OS cells that stably produce GFP-tagged MS2 with a nuclear localization sequence (NLS), as we have done previously^47^. Distinct puncta representing single particles were readily detected in the cytosol (Supplementary Fig. 4A). As with the Riboglow-tagged mRNA, the intensities of the puncta in an ROI follow a Gaussian distribution (Supplementary Fig. 4C), indicating single MS2-GFP tagged ACTB mRNA^7^. (iv) We did not observe appreciable variability in particle size for Riboglow-tagged ACTB or ACTB tagged with 24xMS2-SL (Supplementary Fig. 5), a feature that would be indicative of fluorophore or particle aggregation. We concluded that we are able to detect single mRNAs when tagging the reporter ACTB mRNA with twelve copies of Riboglow.

We developed and validated a pipeline to quantify single ACTB mRNA tagged with 24 copies of the MS2-SL (Supplementary Fig. 4) and took advantage of existing studies to validate our approach. Qualitative inspection of movies revealed different types of movement, as expected (Supplementary Fig. 4): the movement of the majority of particles was non-directional, resembling diffusion (Supplementary Fig. 4B, right side of the field of view). A minority of particles appeared to move in a directional manner, sometimes rapidly changing direction (Supplementary Fig. 4B, left side of the field of view). To classify and quantify particle motions systematically, we employed the FIJI TraJClassifier^53^. Briefly, the TraJClassifier analyzes single particle trajectories as one of four motion types (“directed / active”, “confined”, “subdiffusion” and “normal diffusion”, Supplementary Fig. 4G). We analyzed MS2-tagged mRNA particles in this way yielding traces with a mean duration of 2.75 s (corresponding to 91 frames at a framerate of 33.3 frames per second, Supplementary Fig. 4F). We observed several examples of “directed”, “normal diffusion” and “subdiffusion” motion (Supplementary Fig. 4D, E). As others have reported^3,11^, we saw directed motion of ACTB mRNA only for a minority of traces (∼15%, Supplementary Fig. 4G). The particle velocity for directed motion was 0.92 ± 0.27 µm/s (Supplementary Fig. 4D), well within the range reported in the literature for directed motion of MS2-tagged mRNA^3,9–11^ (Supplementary Fig. 1A, Supplementary Table 1). We further inspected non-directional movement classified as either “normal diffusion” or “subdiffusion” by the TraJClassifier^53^. The diffusion coefficient we observed for normal diffusion was 0.09 ± 0.03 µm^2^/s, comparable with values observed in the literature^3,7–9,11,12,48^ (Supplementary Fig. 1B, Supplementary Table 1). Traces labeled as “subdiffusion” by the TraJClassifier describe restricted movement^53^, resembling a behavior called “corralled” by Fusco et al^3^. Indeed, as Fusco et al^3^ (Supplementary Fig. 1B), we observed “subdiffusion” behavior with a diffusion coefficient of 0.007 ± 0.005 µm^2^/s, vs. faster normal diffusion (0.09 ± 0.03 µm^2^/s, Supplementary Fig. 4E). The strong resemblance of MS2-SL tagged ACTB mRNA dynamics in our hands vs. previous measurements validates our image acquisition, particle detection, tracking and classification approach to accurately detect and quantify movement of single mRNAs.

Having established a pipeline to detect and quantify movement of single ACTB mRNAs in our hands, we asked whether Riboglow-tagging is compatible with single mRNA detection. As for 24x MS2-SL-tagged mRNA, we tracked particles over several seconds and observed random movement resembling diffusion (Fig. 2A, blue trace), or movement in a directional manner with sudden changes in direction (Fig. 2A, red trace). With the constraint that the minimal track length was set as 30 frames, we detected tracks with a mean length of 68 frames (corresponding to 2.0 s), and the longest track was 281 frames (8.4 s, Fig. 2E). All tracks generated with Trackmate were categorized by the TraJClassifier^53^. Directed movement was detected for a minority of traces (6 ± 3%, average and standard deviation between independent experiments) (Fig. 2B). The mean velocity for “directed” tracks of 1.2 ± 0.5 µm/s (Fig. 2C) was comparable with the number for 24x MS2-SL-tagged ACTB mRNA (0.92 ± 0.27 µm/s, Supplementary Fig. 3D) and values observed in the literature^3,7–9,11,12,48^ (Supplementary Fig. 1B, Supplementary Table 1). Particles classified as “normal diffusion” moved with an apparent diffusion coefficient of 0.09 ± 0.03 µm^2^/s (Fig. 2D), comparable with the value we observed for 24x MS2-SL-tagged ACTB (0.09 ± 0.03 µm^2^/s, Supplementary Fig. 4E). Similarly, the diffusion coefficient for traces classified as “subdiffusion” was reduced (0.0029 ± 0.0004 µm^2^/s, Fig. 2D), as for 24x MS2-SL (0.007 ± 0.005 µm^2^/s, Supplementary Fig. 4E). These numbers and trends are consistent with observations in the literature^3,7–9,11,12,48^ (Supplementary Fig. 1B, Supplementary Table 1). We concluded that the Riboglow platform is capable of detecting single mRNAs in live cells and those mRNAs can be tracked over several seconds.

**Figure 2:**
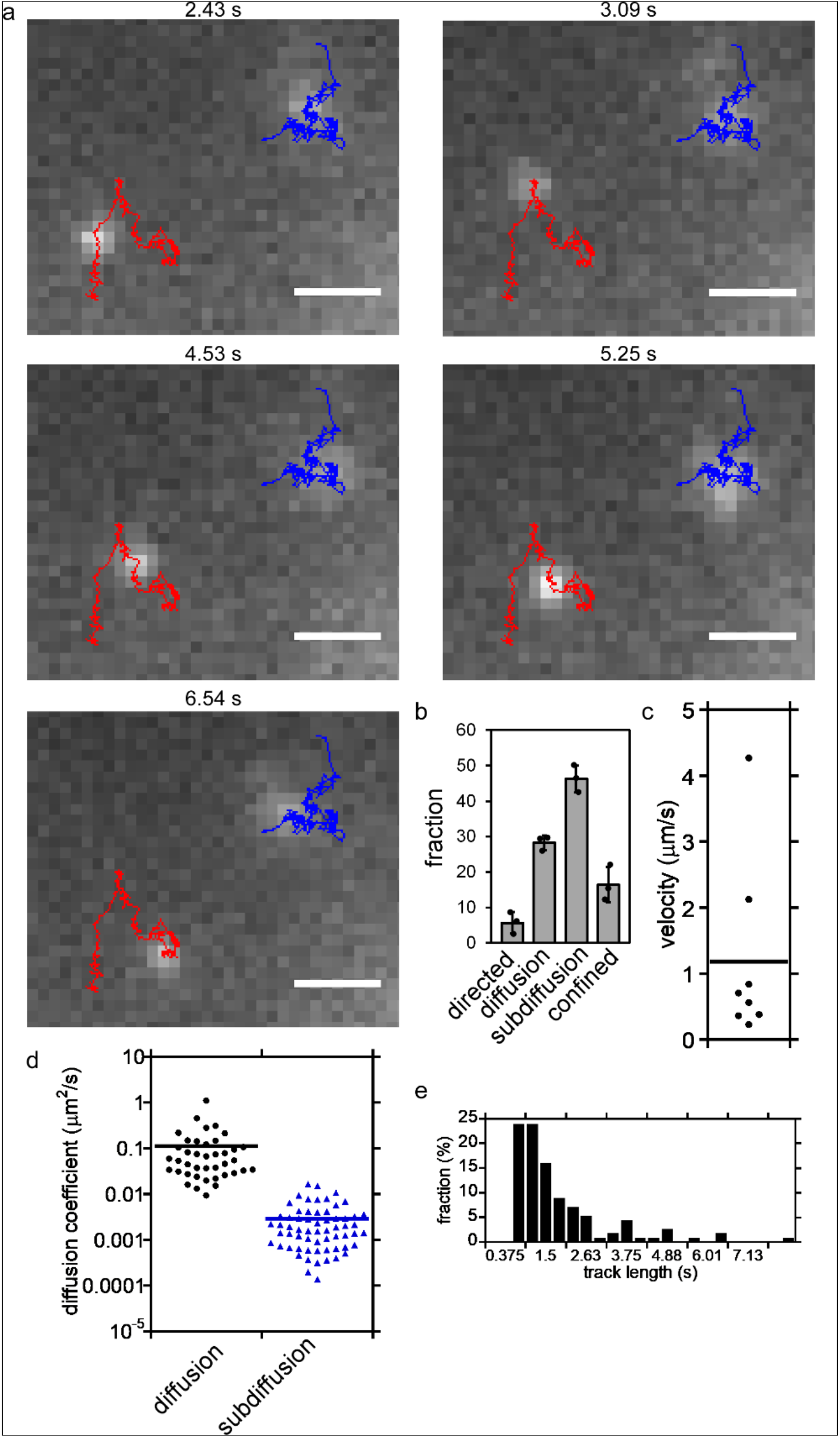
Quantification of single mRNA dynamics with Riboglow-ATTO 590. U2-OS cells were transfected with a plasmid encoding twelve Riboglow-tagged ACTB mRNA, bead loaded with Cbl-4xGly-ATTO 590 and imaged live. (A) Traces of single mRNA particles were generated using the FIJI Trackmate plugin^52^. Shown are representative traces that were classified as “directed” (red, left) and “subdiffusion” (blue, right) with the TraJClassifier^53^. The movie was acquired with 30 ms exposure and a frame rate of 33.3 frames per second. Scale bar = 1 μm. See also Movie 4. (B) Distribution of particle classifications using the FIJI TraJClassifier^53^ for the entire ACTB-12xRiboglow dataset (error bars are STD from three independent experiments). (C) Distribution of velocity for particles classified as moving by “directed movement”^53^. (D) Distribution of diffusion coefficients for particle movements classified as “diffusion” or “subdiffusion” using the FIJI TraJClassifier^53^. (E) Distribution of trace length for particles detected and tracked (113 tracks, mean length = 67.6 frames, corresponding to 2.0 s at a frame rate of 33.3 frames per second). Data from three independent experiments, 6 cells/ 18 ROIs across all experiments.

We sought to explore whether the size of the Riboglow tag can be shortened by reducing the number of tag repeats, as smaller tags are less likely to perturb function of tagged proteins and RNAs. While tagging ACTB mRNA with 12 copies of Riboglow only adds a maximum of 470 kDa to the mRNA, vs. ∼2,500 kDa added when all 24 SLs are bound to a GFP-MS2 dimer in the 24xMS2-SL tag (Supplementary Table 2), we wondered whether a further reduction in tag size may be possible without compromising single mRNA detection. We reduced the number of Riboglow repeats fused to ACTB mRNA to eight copies. As we have demonstrated before^47^, when two plasmids are co-transfected in our hands, the chances of double transfection (e.g. the cell receiving both plasmids) are >90%. We co-transfected a plasmid encoding ACTB tagged with eight copies of the Riboglow tag with a blue marker (NLS-TagBFP), and bead loaded the Cbl-4xGly-ATTO 590 probe. When we only analyzed cells positive for the blue nuclear co-transfection marker, no distinct puncta were visible (Movie 5, compare with Movie 1). Likewise, puncta could not routinely be detected via the Trackmate^52^ algorithm (Fig. 3A). In a small subset of ROIs, occasional puncta were visible and could be detected with Trackmate using our constraints (a radius of 0.5 µm, etc, see Methods), but these puncta were not detectable in subsequent frames and hence we were unable to track them (Fig. 3B). We concluded that robust detection and tracking of single ACTB mRNA molecules with Riboglow requires 12 copies of Riboglow, although a reduction in tag length is likely feasible with improvement of key features of the Riboglow platform, namely fluorescence contrast between the free and bound Cbl-fluorophore probe.

**Figure 3:**
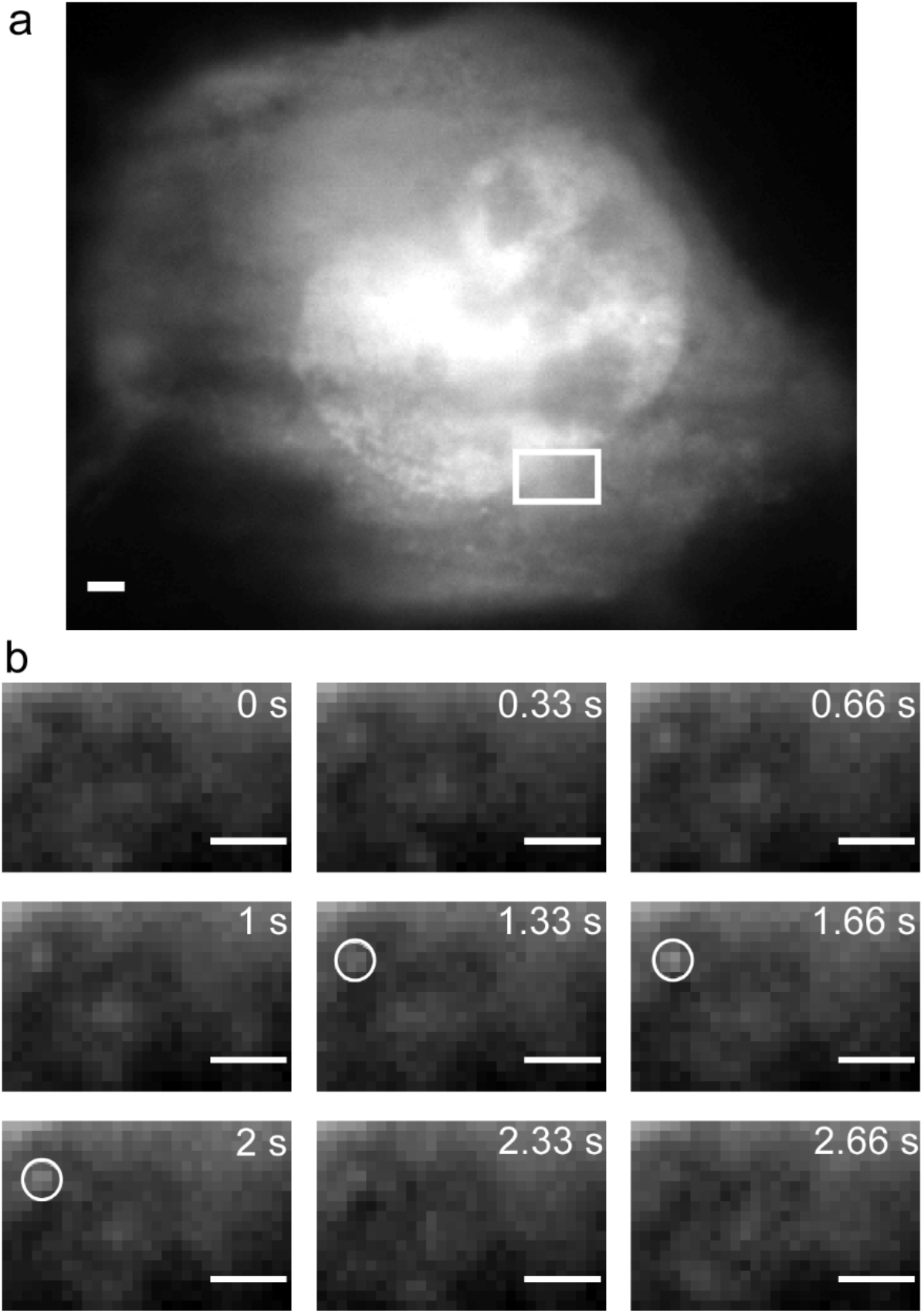
Single particle tracking requires more than 8 copies of Riboglow. A plasmid encoding ACTB tagged with 8 copies of the Riboglow tag “A” was transfected in U2-OS cells together with a transfection marker. The probe Cbl-4xGly-ATTO590 was bead loaded into cells 24 h post transfection. (A) Cells that were positive for the transfection marker and bead loading were interrogated further. No distinct puncta are visible. Scale bar = 2 µm. (B) Insert with a white box in (A) is shown for several consecutive frames. Rarely, distinct puncta are visible and detectable with the Trackmate^52^ algorithm (shown as white circles). Puncta could not be tracked across time points, as illustrated here where a puncta was detected in only 3 consecutive frames (representative example from 2 experiments, 7 cells). Scale bar = 1 µm.

The recent rapid advances in visualizing and quantifying the mRNA lifecycle on the single molecule level using a combination of fluorescent tools prompted us to assess the possibility of monitoring mRNA translation with Riboglow-labeled mRNA. We adapted a translation assay previously developed in the Stasevich lab^22^, where a gene encoding KDM5B has an N-terminal ‘spaghetti monster’ (SM) tag that includes 10 FLAG tag copies to visualize the nascent protein, and the 3’ end includes the MS2-SL mRNA tag. Our KDM5B reporter construct also has the N-terminal SM tag, and we used 12 copies of Riboglow instead of the MS2-tag in the 3’ UTR for mRNA tagging (Supplementary Fig. 2C). The design of this construct allows for detection of the nascent protein with an anti-FLAG antibody fragment (Fab), labeled with green fluorescent Alexa488, as established previously^22,26^. Our reporter gene allows for correct protein translation, confirmed by accumulation of SM-KDM5B labeled with green Fab in cell nuclei (Supplementary Fig. 6). When we loaded the green Fab together with the Riboglow probe Cbl-4xGly-ATTO590 and the SM-KDM5B reporter plasmid DNA, we observed diffraction limited spots where green fluorescence from the nascent protein tag and red fluorescence from the Riboglow-ATTO590 tagged mRNA colocalized in the cytosol (Fig. 4). These spots are consistent with translation sites, for the following reasons. First, we detected co-localization spots around 6 hours post transfection, in line with the timing of translation detection in previous studies^22^. Second, we observed co-movement of these spots, as expected for mRNA translation (Movie 6, Figure 4D, E). At 6-8 hours post transfection, we detected 6 ± 4 spots per cell (6 experiments, 16 cells, 94 total spots) that can be tracked together for at least five consecutive imaging frames (corresponding to > 7 s tracking). Finally, when Puromycin was added to cells with co-moving translation spots, we observed rapid disappearance of the protein signal from the translation sites (Fig. 5B-D). Before Puromycin treatment, 64 ± 14% mRNA spots colocalized with protein signal, vs. 30 ± 12% mRNA spots colocalized with protein signal after Puromycin treatment (3 experiments, 47 mRNA spots total). Puromycin halts translation and releases the elongating polypeptide from the translation site, as used in similar translation assays^22,23^. Together, we established Riboglow as a suitable addition to the growing toolbox of RNA probes to interrogate aspects of the mRNA lifecycle on the single molecule level, including protein translation.

**Figure 4:**
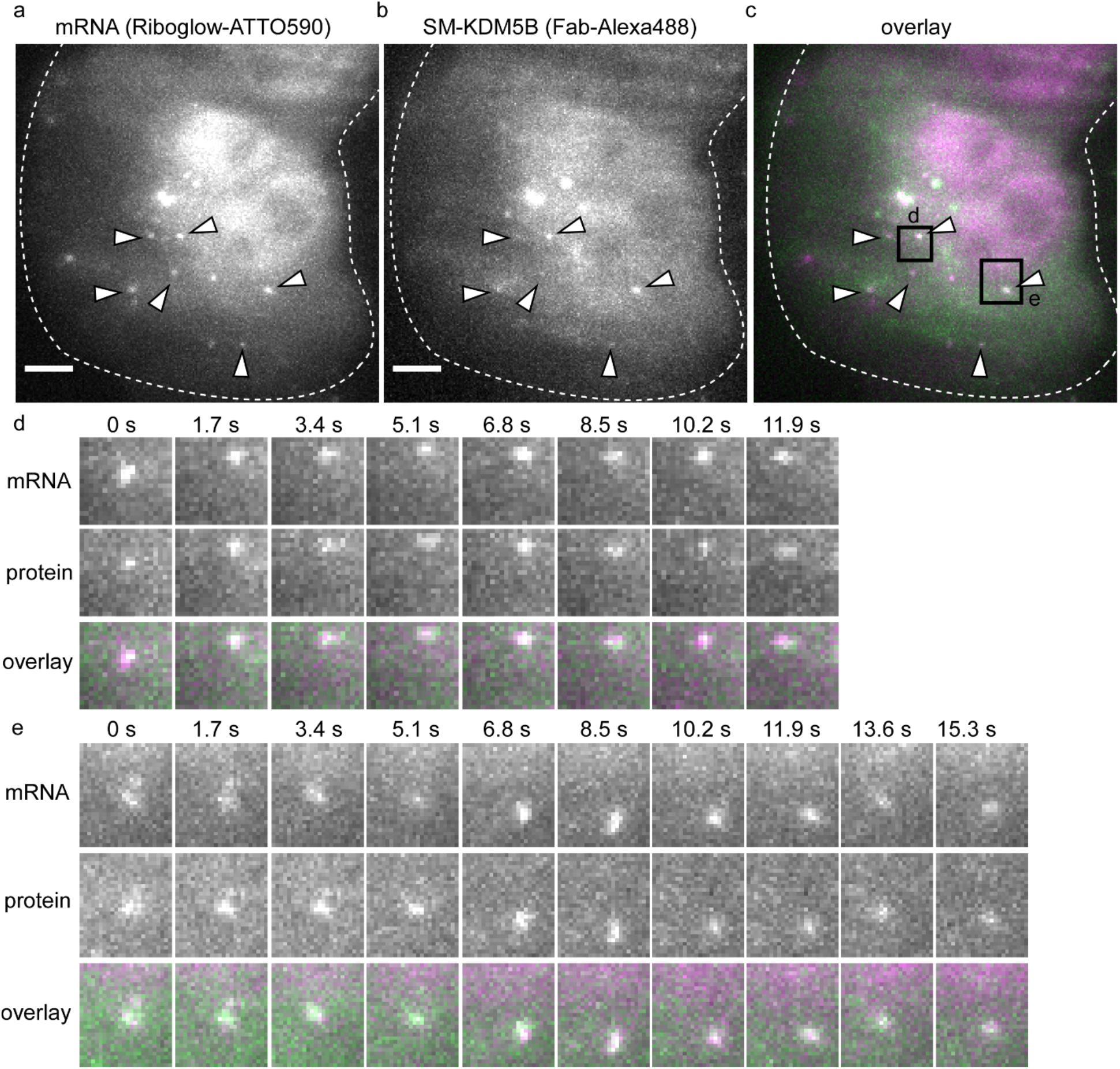
Visualization of mRNA translation with Riboglow-tagged mRNA. U2-OS cells were bead loaded with a plasmid encoding the KDM5B translation assay reporter, together with the Cbl-4xGly-ATTO590 Riboglow probe to label mRNA and Fab-Alexa488 to label the nascent protein. Cells were visualized 6 h after bead loading. Cytosolic spots were detected in both the Riboglow-ATTO590 channel (A) and Fab-Alexa488 channel (B), and these spots co-localize (white arrows). (C) Riboglow-ATTO590 signal is shown in magenta, Fab-Alexa488 is shown in green. (D, E) Tracking of Riboglow-ATTO 590 tagged mRNA and Fab-Alexa488 tagged nascent protein over time (overlay: mRNA in magenta, protein in green). Spots tracked are indicated by black boxes in (C). Maximal intensity projection of six z-stacks (0.5 µm distance per step in z), scale bar = 5 µm.

**Figure 5:**
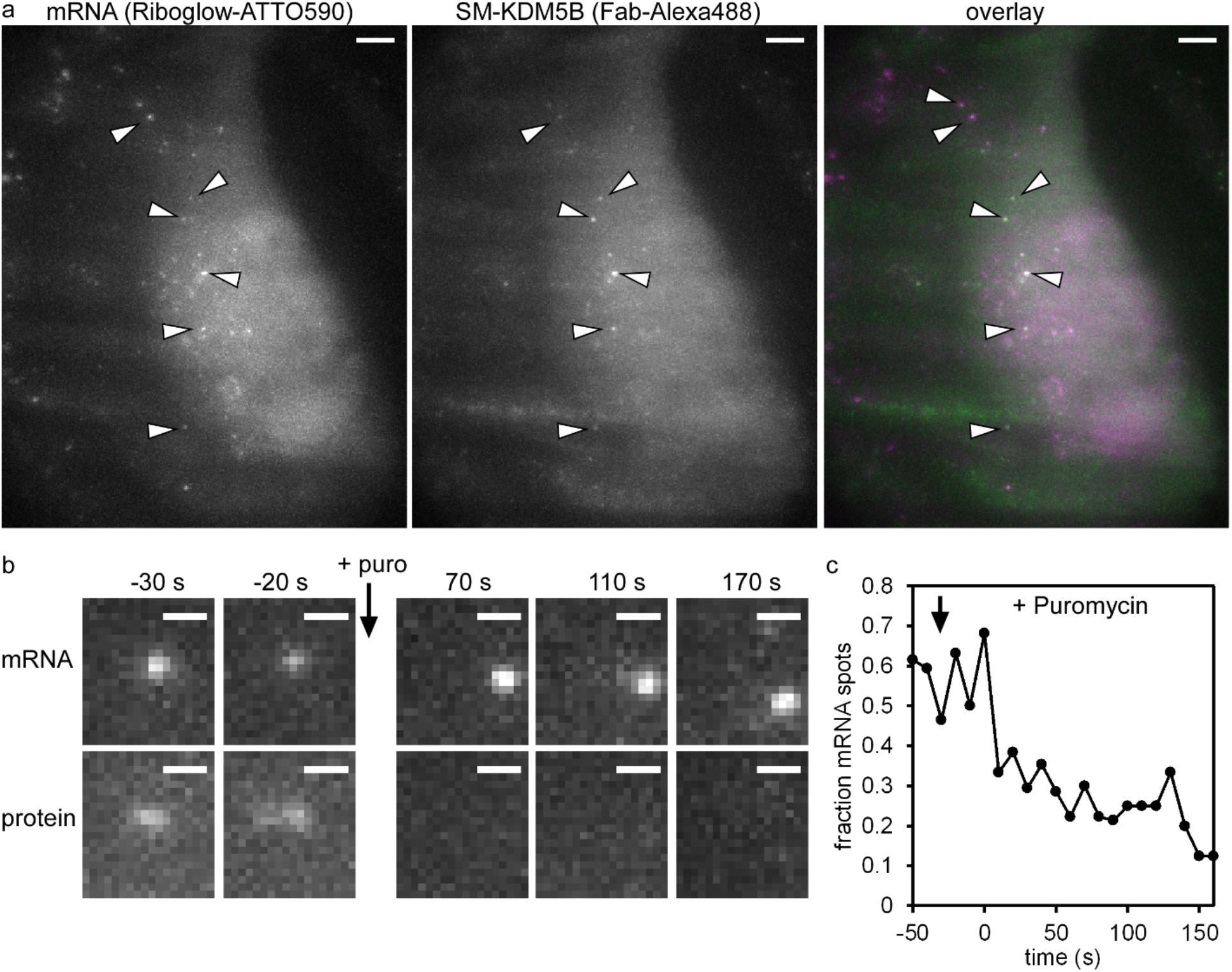
Puromycin-treatment releases the nascent polypeptide from translation sites. The translation assay plasmid DNA encoding for SM-KDM5B was bead loaded together with the Cbl-4xGly-ATTO 590 probe and the green Fab fragment in U2-OS cells. Six hours later, cells positive for Riboglow (Cbl-4xGly-ATTO 590 bound to the tagged mRNA) and SM-KDM5B (where the N-terminal SM was labeled with green fluorescent Fab) were visualized. (A) Fluorescence from Riboglow-ATTO 590, green Fab and an overlay of both is shown where Riboglow-ATTO590 is magenta and Fab is green (colocalization corresponds to white). Several spots where mRNA (labeled with Riboglow-ATTO 590) and nascent protein (labeled with Fab-Alexa 488) colocalize are marked with white arrows, before Puromycin treatment. Maximal intensity projection of 9 z-stacks (0.5 µm distance per step in z), scale bar = 5 µm. Image collection every 10 s. (B) Representative translation spot in the Riboglow-ATTO590 channel (top) and SM-KDM5B channel (bottom) illustrates loss of the SM-KDM5B protein signal from the translation spot upon Puromycin addition. Maximal intensity projection of 9 z-stacks (0.5 µm distance per step in z), scale bar = 1 µm. (C) Quantification of mRNA colocalization with nascent protein. Spots in red (Riboglow-ATTO 590, labeling mRNA) for the cell shown in (A) were counted in each frame, and spots in green (Fab-Alexa 488, labeling nascent protein) were also counted. The fraction of red spots that also display green fluorescence is plotted over time for the cell shown in (A).

## Discussion

In this work we demonstrate Riboglow is a versatile tool capable of detecting and tracking single molecules of RNA in live mammalian cells. We readily detected single mRNA molecules when 12 copies of Riboglow were used and were able to track each mRNA for up to 6 s. Tracking allowed us to characterize and quantify the type of movement. We observed directed movement, diffusion, subdiffusion and confined movement, and quantified parameters for each type (velocity of 1.2 ± 0.5 µm/s for directed tracks, diffusion coefficient of 0.09 ± 0.03 µm^2^/s for diffusive behavior). Our values are consistent with data for single mRNA tagged with 24 copies of MS2-SL, serving as a robust validation for our tool. To demonstrate the potential of Riboglow for multiplexing with other fluorescent reporters, we visualized single Riboglow-tagged mRNA molecules and labeled emerging nascent polypeptide chains at translation sites. Together, we present Riboglow as a robust tool to visualize single mRNAs, and demonstrate its usability for multi-color tracking applications.

Riboglow features several advantageous properties for live RNA labeling. First, Riboglow features organic dyes with photophysical properties ideal for single molecule imaging (high brightness and slow bleaching). Second, we have previously explored exchanging the fluorophore and hence color of Riboglow from red (ATTO 590) to far red (Cy5)^47^ without compromising function. This feature will be desirable for usage of Riboglow in multi-color applications. Third, the size of even a 12 copy tag of Riboglow (471 kDa) is significantly smaller than the conventional 24xMS2 tag (2,527 kDa, assuming 1:1 binding of RNA to dye, Supplementary Table 2). Note that the more recently used MS2-Halo is even larger than MS2-GFP (Halo is 33 kDa, vs. 27 kDa for GFP). It is noteworthy that the smaller overall Riboglow tag vs. the larger 24xMS2 tag did not reveal detectable changes in mRNA dynamics in our hands (Supplementary Fig. 4 vs. Fig. 2). These results suggest that ACTB mRNA mobility in this model system is not likely to be altered by large tags. Regardless, shortening tags for minimal perturbation of the tagged species is advantageous to minimize potential perturbation of the tagged RNA’s function. Lastly, Riboglow is entirely orthogonal to MS2-based mRNA labeling systems, enabling simultaneous labeling of different parts of the same mRNA, or multiple different mRNAs in the same cell. Together, we envision Riboglow as a new member of the growing toolbox for labeling mRNA on the single molecule level.

## Online Methods

### Cloning and mammalian cell biology

A construct to tag ACTB with 12 copies of Riboglow was derived from ACTB-A(4x), described in ref.^47^. Briefly, the entire region of the four-repeat A sequence was PCR amplified with either *Pst*I or *Xba*I overhangs. Plasmid ACTB-A(4x) has unique *Pst*I and *Xba*I sites flanking the A repeat, and the 8mer and 12mer A repeat was generated by sequential ligation using standard molecular cloning protocols (Supplementary Fig. 1). Relevant DNA coding sequences are listed in Supplementary Table 3. All constructs were sequence verified.

U2-OS cells were maintained in Dulbecco’s modified eagle medium (DMEM, Gibco) supplemented with 10% FBS (Gibco) at 37 °C/5% CO_2_. For imaging experiments, U2-OS cells were seeded at 0.2×10^6^ cells in homemade imaging dishes (described in ref.^47^). One day after seeding, cells were chemically transfected with 2 μg plasmid DNA (1 μg of the ACTB-RNA tag fusion mixed with 1 μg of a transfection marker, pNLS-TagBFP, described in ref.^47^) using the TransIT transfection system following manufacturer recommendations (Mirus). On the day of the imaging experiment, 3 μL of a 50 μM stock of Cbl-4xGly-ATTO590 (described in ref.^47^) were bead loaded^54^ per imaging dish, as described in ref.^47^.

### Fab generation and dye-conjugation

Fab generation was done using the Pierce mouse IgG1 preparation kit (Thermo Fisher Scientific) per the manufacturer’s instructions. Briefly, beads conjugated with ficin were incubated in 25 mM cysteine to digest FLAG (Wako, 012-22384 Anti DYKDDDDK mouse IgG_2b_ monoclonal) antibodies to generate Fab. Fab were separated from the digested Fc region using a NAb Protein A column (Thermo Scientific, product # 1860592). Fab were concentrated to ∼1 mg/ml and conjugated to Alexa Fluor 488 (A488). A488 tetrafluorophenyl ester (Invitrogen) was suspended in DMSO and stored at −20°C. 100 µg of Fab were mixed with 10 µL of 1M NaHCO_3_, to a final volume of 100 µL. 5 μl of A488 was added to this 100 µL mixture and incubated for 2 hours at room temperature with end-over-end rotation. The dye conjugated Fab were eluted from a PBS equilibrated PD-mini G-25 desalting column (GE Healthcare) to remove unconjugated dye. Dye conjugated Fabs then were concentrated in an Ultrafree 0.5 filter (10k-cut off; Millipore) to 1 mg/ml. This conjugation and concentration process was repeated on occasion to ensure a degree of labeling close to one. The ratio of Fab:dye, *A*_*rat*_, was determined using the absorbance at 280 and 495 nm, the extinction coefficient of IgG at 280 nm, ε_*IgG*_, the extinction coefficient of the dye, ε_*dye*_, provided by the manufacturer, and the dye correction factor at 280 nm, *CF*, provided by the manufacturer. The degree of labeling, *DOL*, was calculated with the following formula:

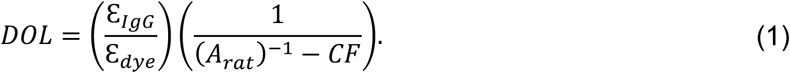

Only Fab calculated with a *DOL* ∼1 were used in experiments.

### Single molecule live imaging

All single molecule live cell imaging was carried out on a Nikon N-STORM microscope, equipped with a TIRF illuminator, 405 nm, 488 nm, 561 nm, and 647 nm laser lines, an environmental chamber to control humidity and temperature, an ORCA-Flash4.0 V2 C11440 sCMOS camera (Hamamatsu) (unless otherwise indicated), a 100x oil-immersion objective (Nikon, NA = 1.49), two filter wheels, and the appropriate filter sets. Cells were cultured on homemade imaging dishes as described previously^47^. Unless otherwise noted, cells were transfected with a plasmid to produce tagged mRNA 24 h prior to imaging. For Riboglow imaging, 3 µL of a 5 µM stock of the Cbl-fluorophore probe Cbl-4xGly-ATTO 590 were bead loaded immediately prior to imaging as described previously^47^, unless otherwise indicated. Cells were washed 3 times with PBS, and medium was replaced with FluoroBrite medium (Gibco). Imaging was performed under HILO conditions^49^ at 37°C and 5% CO_2_. Details on imaging acquisition conditions are summarized in Supplementary Table 4.

### Particle detection and tracking

Particles were detected using the FIJI plugin TrackMate^52^ (version 3.8.0). Movies were divided into small regions of interest (ROI), as illustrated in Fig. 2. Because a low density of particles is optimal to allow for tracking of the same particles across frames, regions towards the edge of the cell were chosen frequently as ROIs. This was done for 24xMS2-SL tagged mRNA and for Riboglow-tagged mRNA, such that any resulting bias affects both mRNA tags. Detection in TrackMate was performed with the LoG detector, an estimated blob size of 0.5 micron, and a Signal/Noise ratio of 0.5-1.0 was used as a filter for each ROI (adjusted manually). For tracking, the simple LAP tracker was used. Linking maximum distance and Gap-closing maximum distance were both set to 0.5 micron, and Gap-closing maximum frame gap was set as 2. Resulting tracks were further filtered to only include tracks with 30 frames per spot. Spot detection, tracking and filters were manually validated by inspecting each ROI movie. Classification of tracks was done by the TraJClassifier^53^. The minimal track length was set to 30 frames. Results from tracking are reported as mean values ± standard error, unless otherwise indicated.

### Translation assay

U2-OS cells seeded on home-made imaging dishes as described above were bead loaded with a mix of 1 µg plasmid DNA encoding for the SM-KDM5B translation reporter, 3 µL of a 5 µM stock of the Cbl-fluorophore probe Cbl-4xGly-ATTO 590 and 1 µL of Alexa488-labeled Fab to bind N-terminal FLAG tags in the reporter. After 6 – 8 hours, cells were washed 3 times with PBS, and medium was replaced with FluoroBrite medium (Gibco). Imaging was performed under HILO conditions^49^ at 37°C and 5% CO_2_ using an Andor Ixon Ultra DU897U - C50 EMCCD camera. Details on imaging acquisition conditions are summarized in Supplementary Table 4. When indicated, puromycin was added during the image acquisition at a final concentration of 50 µg/mL.

## Acknowledgements

We acknowledge support from the National Institute of Health (K99 GM127752 to E.B., R01 GM133184 to A.E.P. and R.T.B., DP1 GM114863 to A.E.P., R01 GM073850 to R.T.B., 5R35 GM119728 to T.J.S.).

## Author contributions

E.B., R.T.B. and A.E.P. conceptualized the study. A.P and E.B. designed experiments. E.B. performed experiments, and analyzed data with input from all authors. K.L. labeled and purified fluorescent Fab fragment. T.J.S. designed translation assay and assisted with implementation of the assay. E.B. wrote the manuscript with edits from all authors.

## Data availability

The datasets generated during and/or analyzed during the current study are available from the corresponding authors on reasonable request.

