## Supporting information (Figures and Tables) for "Detection and quantification of single mRNA dynamics with the Riboglow fluorescent RNA tag"

Supplementary Figure 1: Literature values for mRNA movement

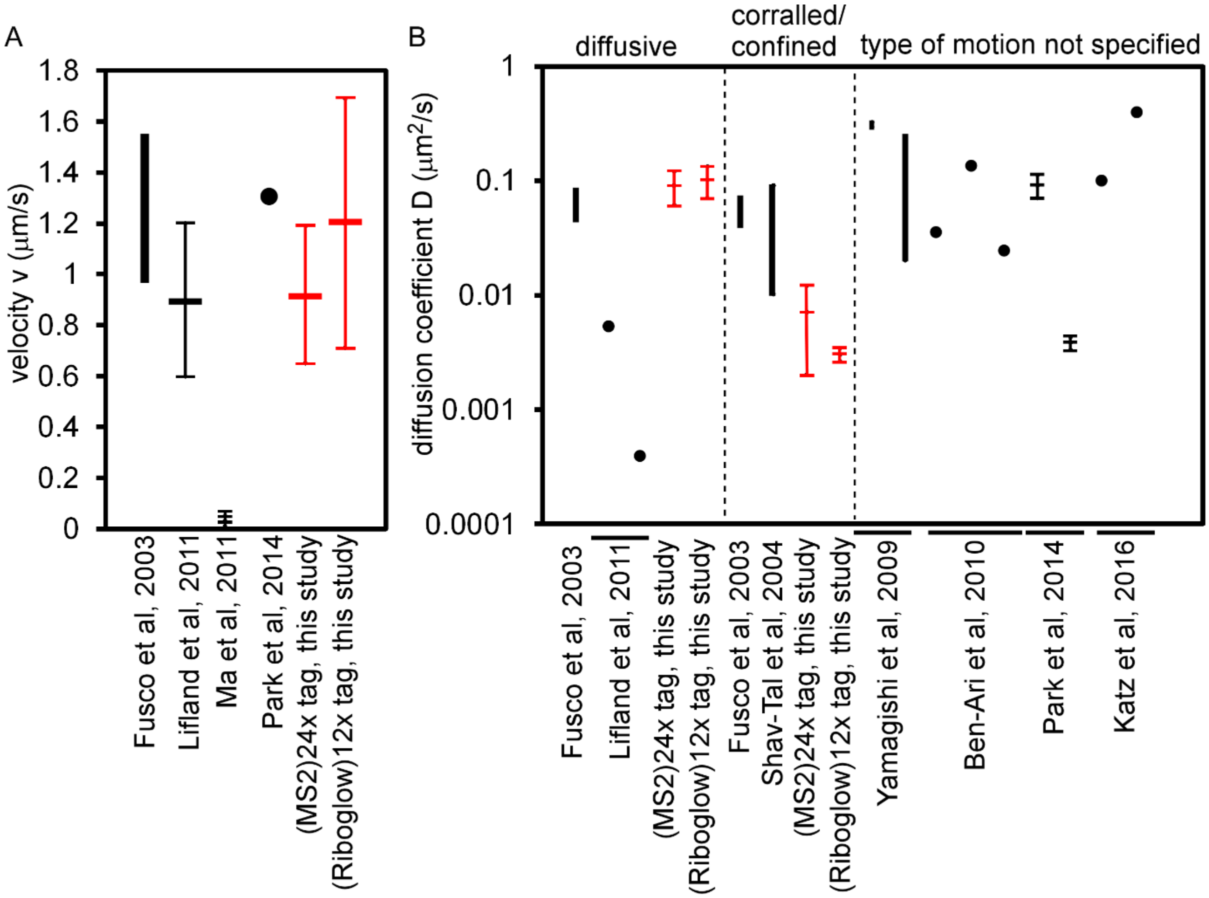


Summary of literature values for single mRNA movement in live cells. Figure demonstrates (A) velocity (v) reported for directed movement, and (B) diffusion coefficient D reported for non-directed movement. The type of motion is indicated at the top of the plot, if specified. Black bar indicates range of values reported. See also Supplementary Table 1 for more details of literature studies. Error bars for data reported from this study represent the standard error.

Supplementary Figure 2: Plasmid maps

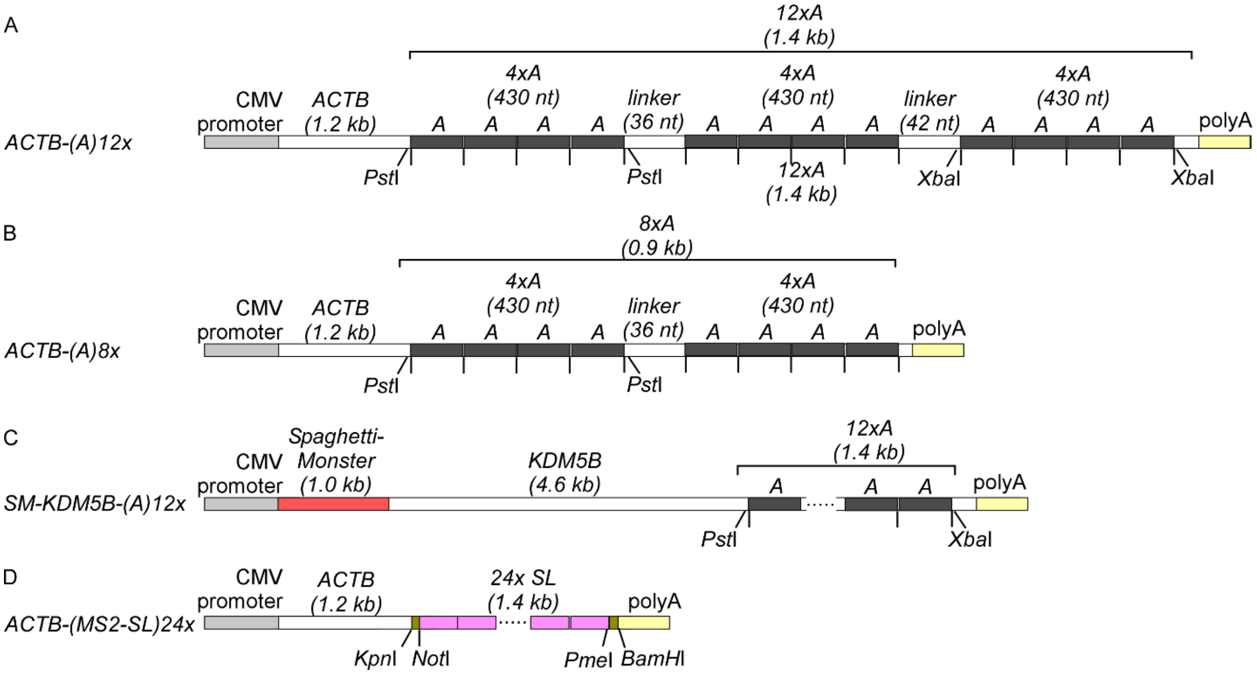


Maps of plasmids used on this study (not drawn to scale) (see also Supplementary Table 3 for tag sequences). (A) 12 copies of the Riboglow RNA-tag referred to as "A" in ref.[47] were fused to the ACTB gene, resulting in construct ACTB-(A)12x. (B) 8 copies of the Riboglow-RNA tag referred to as "A" were fused to the ACTB gene, resulting in construct ACTB-(A)8x. (C) The KDM5B gene was N-terminally tagged with the 10xFlag-tag marker termed spaghetti-monster in Ref[55] and tagged with 12x Riboglow as in (A) in the 3'UTR. (D) The ACTB gene was tagged with 24 copies of the MS2SL sequence as described in ref[47].

Supplementary Figure 3: Particle intensity distribution in movie with ACTB mRNA tagged with 12 copies of Riboglow for region of interest (ROI) fit with two Gaussian distributions

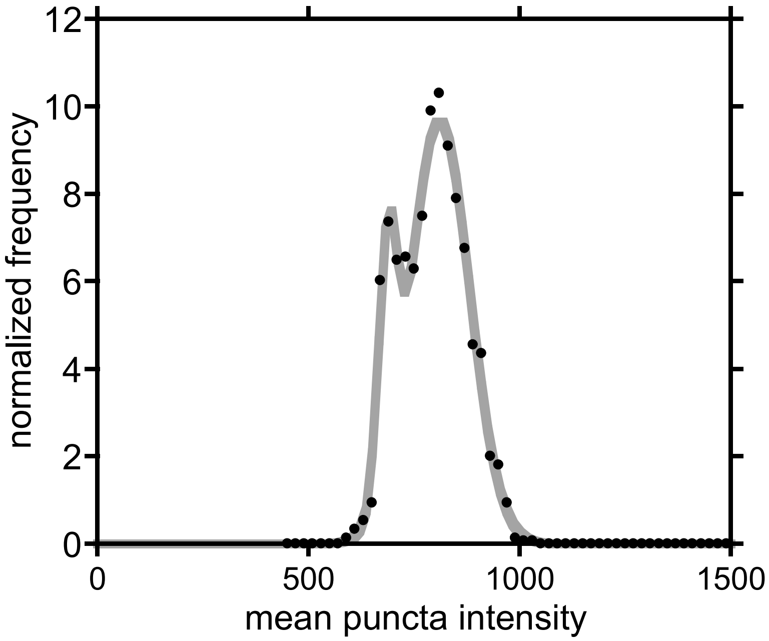


Some regions of interest (ROI) in movies where fluorescent particles are tracked display particle intensity distributions that can be described with a multiple Gaussian fit. N = 1496 puncta, R = 1.0.

Supplementary Figure 4:  Detection of single ACTB mRNA with the MS2 system

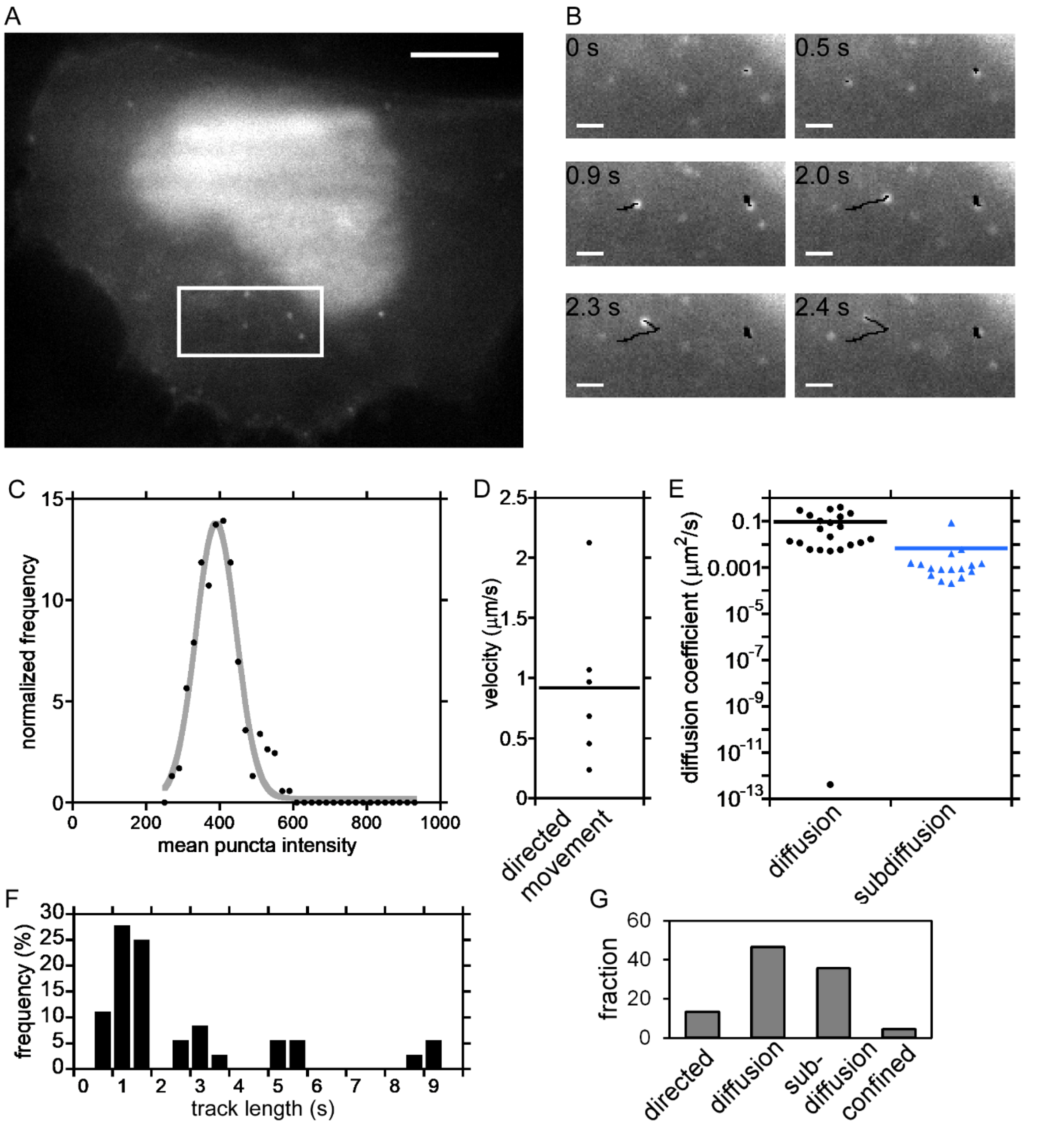


U2-OS cells that stably produce NLS-MS2-GFP were transiently transfected with a plasmid producing ACTB mRNA tagged with 24 copies of the MS2 stem loop (SL) sequence and imaged 24 h later.  (A) Shown is a representative cell where distinct particles were detected in the green fluorescence channel, representing single mRNAs.  Scale bar = 5 μm.  White box: panel (B) zoom in.  (B) Time frames of zoomed in region (white box in panel (a)) reveal dynamics of mRNA movement.  Movies were acquired at a frame rate of 33.3 frames per second, and select frames are shown. Particles were detected and tracked using the FIJI tracking plugin TrackMate [52]. Shown are two tracked particles; the black lines represent traces over time. (C) Particles detected using TrackMate have fluorescence intensities that follow a Gaussian distribution. Shown are all particles detected in a representative region of interest (ROI) used for tracking analysis over time (n = 532 particles). The intensity distribution was fit with a Gaussian distribution (R = 0.98). The type of movement for each tracked particle from the TrackMate analysis was classified using the FIJI TraJClassifier[53]. (D) Distribution of velocity for particles classified as moving by "directed movement"[53]. (E) Distribution of diffusion coefficients for particle movements classified as "diffusion" or "subdiffusion" using the FIJI TraJClassifier[53]. (F) Length distribution of tracked particle traces (36 traces total, mean: 2.75 s / 90.58 frames at a frame rate of 33.3 frames per second). (G) Distribution of particle classifications using the FIJI TraJClassifier[53] for the entire ACTB-24xMS2-SL dataset. Data from two experiments, three cells, 5 ROIs.

Supplementary Figure 5: Particle size estimates for spot detection

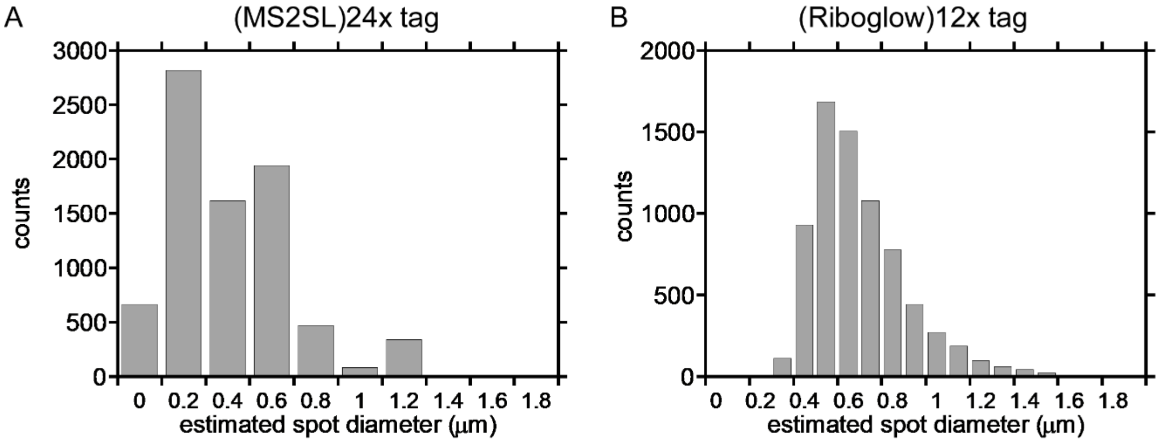


Estimate of diameter for spots detected by TrackMate. (A) Distribution of estimated diameter for all spots for analysis of (MS2)24x-tagged ACTB mRNA (7935 spots total). (B) Representative distribution of estimated diameter for spots of (Riboglow)12x-tagged ACTB mRNA (7242 spots total).

Supplementary Figure 6: KDM5B in the translation reporter construct localizes to the nucleus

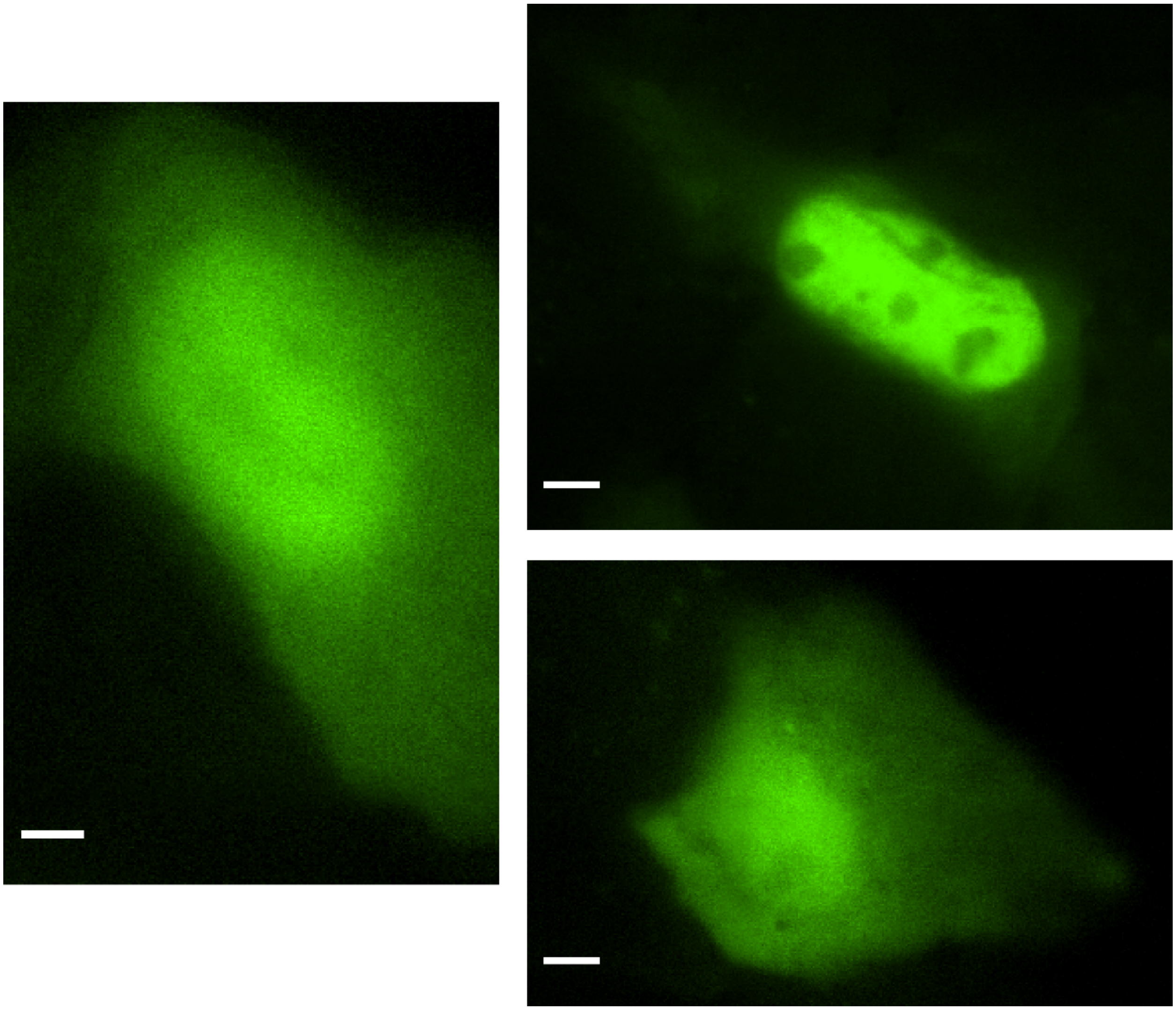

U2-OS cells were bead loaded with a mix of the SM-KDM5B translation reporter plasmid DNA and a green fluorescent Fab fragment to label the N-terminus of the resulting SM-KDM5B protein. Cells were imaged 7 h post bead loading of DNA, Fab fragment, and Riboglow probe (cells from 2 separate experiments are shown). Scale bar = 5 μm.

Supplementary Table 1: Literature values for live mRNA dynamic measurements (relevant studies)

D = diffusion coefficient; v = velocity; SPT = single particle tracking; FRAP = fluorescence recovery after photobleaching, MS2-SL = MS2 stem loop

| **Key experimental conditions** | **Quantification of movement** | **Reference** |
|---|---|---|
| lacZ tagged with 24x MS2-SL, in COS cells | Directed movement (2-5%): v = 1 – 1.5 μm/s; diffusional movement (D = 0.08 – 0.045 $\mu m^2/s$); corralled movement (D = 0.07 – 0.0413 $\mu m^2/s$) | 3 |
| YFP tagged with 24x MS2-SL, stably integrated in U2OS cells, tracking of nuclear movement | D = 0.01 – 0.09 $\mu m^2/s$ (nucleus, SPT) (simple diffusion, corralled diffusion); D = 0.09 ± 0.0006 $\mu m^2/s$ (nucleus, FRAP) | 48 |
| Beta-actin tagged with 24x MS2-SL, in chicken embryo fibroblast (CEF) cells | D = 0.32 – 0.29 $\mu m^2/s$ (leading edge of the cell); D = 0.02 – 0.24 $\mu m^2/s$ (perinuclear region) | 7 |
| Chicken beta-actin (with its 3'UTR zipcode), tagged with CFP (N-terminus) and 24x MS2-SL, stably integrated in U2OS cells | D = 0.035 $\mu m^2/s$ (cytoplasm, SPT); D = 0.13 $\mu m^2/s$ (cytoplasm, FRAP), D = 0.024 $\mu m^2/s$ (nucleoplasm, SPT) | 8 |
| Endogenous beta-actin, labeled with MTRIPs (multiply-labeled tetravalent imaging probes), in A549 cells (epithelial cell line) | SPT: active, processive motion (v = 0.9 ± 0.3 μm/s) and passive, diffusive motion (0.0053 $\mu m^2/s$ in A549 cells, 0.0004 $\mu m^2/s$ in fibroblasts) | 9 |

| | | |
|---|---|---|
| and human dermal fibroblasts (primary) | | |
| Beta-actin tagged with 24x MS2-SL, in primary rat cortical neurons | v = 0.0294 +/- 0.0215 μm/s | 10 |
| Beta-actin tagged with 24x MS2-SL, in mice. Imaging in primary cells from those mice. | SPT: In primary fibroblasts: D = 0.09 ± 0.02 $\mu m^2/s$, ~1% directed movement; in primary neurons: D = 3.8 (± 0.5) x $10^{-3}$ $\mu m^2/s$, ~10% active transport (v = 1.3 μm/s) | 11 |
| Endogenous beta-actin tagged with 24x MS2-SL, fibroblast cells derived from transgenic mouse | D = 0.1 (slow) - 0.4 (fast) $\mu m^2/s$, shift to fast state upon puromycin treatment (ribosome release) | 12 |

Supplementary Table 2: Size comparison of RNA tags

| Name of tag component | Size estimate (kDa) |
|---|---|
| 12x Riboglow RNA tag (A variant[47]) | 442 kDa |
| 8x Riboglow RNA tag (A variant[47]) | 290 kDa |
| Cbl-4xGly-ATTO590 | 2.4 kDa |
| 12x Riboglow RNA tag bound to 12 Cbl-4xGly-ATTO590 probes | 471 kDa |
| 8x Riboglow RNA tag bound to 8 Cbl-4xGly-ATTO590 probes | 309 kDa |
| 24x MS2-SL RNA tag | 415 kDa |
| MS2-GFP homo dimer | 88 kDa |
| 24x MS2-SL RNA tag bound to 24 MS2-GFP homo dimers | 2,527 kDa |

Supplementary Table 3: Sequences for relevant plasmid regions

| | |
|---|---|
| Riboglow DNA sequence (1 copy, A in ref[47]) | CCT AAA AGC GTA GTG GGA AAG TGA CGT GAA ATT CGT CCA GAT<br>TAC TTG ATA CGG TTA TAC TCC GAA TGC CAC CTA GGC CAT ACA<br>ACG AGC AAG GAG ACT CA |
| 8xRiboglow sequence (Riboglow sequence in *italic*, linkers in **bold**) | *CCT AAA AGC GTA GTG GGA AAG TGA CGT GAA ATT CGT CCA GAT*<br>*TAC TTG ATA CGG TTA TAC TCC GAA TGC CAC CTA GGC CAT ACA*<br>*ACG AGC AAG GAG ACT CA***G GTA CCG G***CC TAA AAG CGT AGT GGG*<br>*AAA GTG ACG TGA AAT TCG TCC AGA TTA CTT GAT ACG GTT ATA*<br>*CTC CGA ATG CCA CCT AGG CCA TAC AAC GAG CAA GGA GAC TCA*<br>**GGA TCC GG***C CTA AAA GCG TAG TGG GAA AGT GAC GTG AAA TTC*<br>*GTC CAG ATT ACT TGA TAC GGT TAT ACT CCG AAT GCC ACC TAG*<br>*GCC ATA CAA CGA GCA AGG AGA CTC A***GG ATC CGG** *CCT AAA AGC*<br>*GTA GTG GGA AAG TGA CGT GAA ATT CGT CCA GAT TAC TTG ATA*<br>*CGG TTA TAC TCC GAA TGC CAC CTA GGC CAT ACA ACG AGC AAG*<br>*GAG ACT CA***G GAT CCA CCG GAT CTA GCT GCA GTC GAC GGT ACC**<br>**GG***C CTA AAA GCG TAG TGG GAA AGT GAC GTG AAA TTC GTC CAG*<br>*ATT ACT TGA TAC GGT TAT ACT CCG AAT GCC ACC TAG GCC ATA*<br>*CAA CGA GCA AGG AGA CTC A***GG TAC CGG** *CCT AAA AGC GTA GTG*<br>*GGA AAG TGA CGT GAA ATT CGT CCA GAT TAC TTG ATA CGG TTA*<br>*TAC TCC GAA TGC CAC CTA GGC CAT ACA ACG AGC AAG GAG ACT*<br>*CA***G GAT CCG G***CC TAA AAG CGT AGT GGG AAA GTG ACG TGA AAT*<br>*TCG TCC AGA TTA CTT GAT ACG GTT ATA CTC CGA ATG CCA CCT*<br>*AGG CCA TAC AAC GAG CAA GGA GAC TCA* **GGA TCC GG***C CTA AAA*<br>*GCG TAG TGG GAA AGT GAC GTG AAA TTC GTC CAG ATT ACT TGA* |

| | |
|---|---|
| | *TAC GGT TAT ACT CCG AAT GCC ACC TAG GCC ATA CAA CGA GCA AGG AGA CTC A***GG ATC C** |
| 12xRiboglow sequence (Riboglow sequence in *italic*, linkers in **bold**) | *CCT AAA AGC GTA GTG GGA AAG TGA CGT GAA ATT CGT CCA GAT TAC TTG ATA CGG TTA TAC TCC GAA TGC CAC CTA GGC CAT ACA ACG AGC AAG GAG ACT CA***G GTA CCG G***CC TAA AAG CGT AGT GGG AAA GTG ACG TGA AAT TCG TCC AGA TTA CTT GAT ACG GTT ATA CTC CGA ATG CCA CCT AGG CCA TAC AAC GAG CAA GGA GAC TCA* **GGA TCC GG***C CTA AAA GCG TAG TGG GAA AGT GAC GTG AAA TTC GTC CAG ATT ACT TGA TAC GGT TAT ACT CCG AAT GCC ACC TAG GCC ATA CAA CGA GCA AGG AGA CTC A***GG ATC CGG** *CCT AAA AGC GTA GTG GGA AAG TGA CGT GAA ATT CGT CCA GAT TAC TTG ATA CGG TTA TAC TCC GAA TGC CAC CTA GGC CAT ACA ACG AGC AAG GAG ACT CA***G GAT CCA CCG GAT CTA GCT GCA GTC GAC GGT ACC GG***C CTA AAA GCG TAG TGG GAA AGT GAC GTG AAA TTC GTC CAG ATT ACT TGA TAC GGT TAT ACT CCG AAT GCC ACC TAG GCC ATA CAA CGA GCA AGG AGA CTC A***GG TAC CGG** *CCT AAA AGC GTA GTG GGA AAG TGA CGT GAA ATT CGT CCA GAT TAC TTG ATA CGG TTA TAC TCC GAA TGC CAC CTA GGC CAT ACA ACG AGC AAG GAG ACT CA***G GAT CCG G***CC TAA AAG CGT AGT GGG AAA GTG ACG TGA AAT TCG TCC AGA TTA CTT GAT ACG GTT ATA CTC CGA ATG CCA CCT AGG CCA TAC AAC GAG CAA GGA GAC TCA* **GGA TCC GG***C CTA AAA GCG TAG TGG GAA AGT GAC GTG AAA TTC GTC CAG ATT ACT TGA TAC GGT TAT ACT CCG AAT GCC ACC TAG GCC ATA CAA CGA GCA AGG AGA CTC* **AGG ATC CAC CGA GTC TAG AAG CAT TGC AGT CGA CGG TAC CGG** *CCT AAA AGC GTA GTG GGA AAG TGA CGT GAA ATT* |

| | |
|---|---|
| | *CGT CCA GAT TAC TTG ATA CGG TTA TAC TCC GAA TGC CAC CTA*<br>*GGC CAT ACA ACG AGC AAG GAG ACT CA***G GTA CCG** *GCC TAA AAG*<br>*CGT AGT GGG AAA GTG ACG TGA AAT TCG TCC AGA TTA CTT GAT*<br>*ACG GTT ATA CTC CGA ATG CCA CCT AGG CCA TAC AAC GAG CAA*<br>*GGA GAC TCA* G**GA TCC GG***C CTA AAA GCG TAG TGG GAA AGT GAC*<br>*GTG AAA TTC GTC CAG ATT ACT TGA TAC GGT TAT ACT CCG AAT*<br>*GCC ACC TAG GCC ATA CAA CGA GCA AGG AGA CTC A***GG ATC CGG**<br>*CCT AAA AGC GTA GTG GGA AAG TGA CGT GAA ATT CGT CCA GAT*<br>*TAC TTG ATA CGG TTA TAC TCC GAA TGC CAC CTA GGC CAT ACA*<br>*ACG AGC AAG GAG ACT CA*G GAT CC |

Supplementary Table 4: Image acquisition

| Experiment | Image acquisition details |
|---|---|
| Live imaging of ACTB-(MS2-SL)24x mRNA | 30 ms exposure, no binning or 2x2 binning , 13.3 or 33.3 frames per second, movie length typically 300 frames, 488 nm laser (~15% power) |
| Live imaging of ACTB-(Riboglow)12x mRNA | 30 ms exposure, 2x2 binning, movie acquisition at 512x512 pixel and 600 frames (33.3 frames per second), 561 nm laser (50% power) |
| Live imaging of ACTB-(Riboglow)8x mRNA | 30 ms exposure, 2x2 binning, movie acquisition at 512x512 pixel and 600 frames (33.3 frames per second), 561 nm laser (50% power) |
| Live imaging of ACTB-(Riboglow)12x mRNA, translation assay | 31 ms exposure, no binning, 1 frame per 10 s, 561 nm laser (15% power), 488 nm laser (15% power), image acquired for 11 z stacks at 0.5 μm per stack |

Movie 1: A plasmid encoding ACTB mRNA tagged with 12 copies of Riboglow was transfected in U2-OS cells, followed by bead loading of Cbl-4xGly-ATTO 590 24 h post transfection, as in Figure 1. Movies of live cells reveal rapid movement of red fluorescent puncta. The movie was acquired with 30 ms exposure and a frame rate of 33.3 frames per second. Scale bar = 2 μm.

Movie 2 and 3: Comparison of U2-OS cells that were bead loaded with Cbl-4xGly-ATTO 590, with (movie 2) or without (movie 3) transfection of a plasmid encoding ACTB mRNA tagged with 12 copies of Riboglow 24 h prior. Red fluorescent puncta that rapidly move in cells are only detectable in cells that were transfected. Movies were acquired with 30 ms exposure and a frame rate of 13 frames per second. Scale bar = 10 μm.

Movie 4: Example of single particle tracking for puncta detected in U2-OS cells that were transfected with 12x Riboglow-tagged ACTB and subsequently bead loaded with Cbl-4xGly-ATTO 590. Shown are traces that were classified as "directed" (red) and "subdiffusion" (blue). Select frames of this movie are shown in Fig. 2A. The movie was acquired with 30 ms exposure and a frame rate of 33.3 frames per second. Scale bar = 1 μm.

Movie 5: A plasmid encoding ACTB mRNA tagged with 8 copies of Riboglow was transfected in U2-OS cells, followed by bead loading of Cbl-4xGly-ATTO 590 24 h post transfection, as in Movie 1. Only cells that were transfected, as judged by the presence of the blue nuclear co-transfection marker protein, were chosen for analysis. The movie was acquired with 30 ms exposure and a frame rate of 33.3 frames per second. Scale bar = 2 μm.

Movie 6: U2-OS cells were bead loaded with a plasmid encoding the SM-KDM5B translation assay reporter, together with the Cbl-4xGly-ATTO 590 Riboglow probe to label mRNA and Fab-Alexa488 to mark the nascent protein. Cells were visualized 6 h after bead loading. (A) Fab-

Alexa488 channel (B) Riboglow-ATTO 590 channel, (C) overlay with Riboglow-ATTO590 signal is shown in magenta, Fab-Alexa488 is shown in cyan. Maximal intensity projection of six z-stacks (0.5 μm distance per step in z), scale bar = 5 μm. 0.7 s frame rate, movie acquisition for 11 slices in z, 18 frames total (30 s).
